# The latent load and the population-optimal mutation rate

**DOI:** 10.1101/2023.06.26.546616

**Authors:** Gordon Irlam

**Affiliations:** Los Altos, California, United States

**Keywords:** positive selection, purifying selection, negative selection, genetic load, latent load, substitutional load, lag load, mutational load, drift load, optimal mutation rate

## Abstract

The rarity of positively selected sites may lead to the expectation that they have only a minor effect on total genetic load. A framework for the genetic load associated with positively selected sites in sexual populations is presented and analyzed. This framework defines the latent load as the genetic load associated with positively selected sites that are destined to fix, but for which beneficial alleles have not yet become established. A formula for the latent load is derived that incorporates various real-world complicating factors. In humans, the latent load is estimated to be larger than the mutational load, and much larger than the substitutional load. The germline spontaneous mutation rate that maximizes population mean fitness is predicted within this framework to occur when the mean latent load is approximately equal to the mutational load. This population-optimal mutation rate differs from the minimal mutation rate predicted by purely microevolutionary considerations that focus on individual-level selection. It is hypothesized that, over macroevolutionary timescales, species-level processes such as differential extinction may bias the mutation rates of surviving species toward the population-optimal value. Overall, this framework highlights the importance of positively selected sites to the total genetic load, and suggests a potential role for the latent load in shaping the long-term evolution of the mutation rate.

## Introduction

The goals of this manuscript are twofold: to develop an equation that highlights the importance of the latent load associated with positively selected sites, and to use this equation to understand how aggregate population mean fitness changes with the germline spontaneous mutation rate.

The latent load is defined here as the genetic load associated with sites that could harbor beneficial alleles but at which such alleles have not yet arisen and begun to fix. For humans, the latent load is shown to be larger than the mutational load, and to exceed the substitutional load by more than two orders of magnitude. Despite this, the latent load appears to have rarely been quantitatively considered by population geneticists. Maynard Smith defines the related concept of the “lag load” (Maynard Smith, 1976), but does not provide a corresponding mathematical formulation. Subsequently several authors have mathematically explored the lag load, but have done so through evolution in an abstract phenotypic space (Lynch and Lande, 1993; Chevin, 2013), rather than as a directly quantifiable load. A simple formula for the population mean fitness associated with the latent load is developed that incorporates various real-world factors. The real-world factors considered are conditional selection coefficients, standing variation, epistasis, nearly neutral positively selected sites, clustering of new positively selected sites in time and space, macroevolutionary selection, and the gradual emergence of a beneficial allele’s positive selection coefficient.

Population mean fitness is important because it reflects the expected number of organisms present in the next generation, and is therefore a key contributor to the success of the population or species. An important result derived here is that the mutation rate per base pair that maximizes the long-term geometric-mean population mean fitness is the one for which the mean fitness losses associated with the latent load and the mutational load are approximately equal. This mutation rate will be termed the population-optimal mutation rate. This finding is significant because it is a mutation rate that is quite different from the as-small-as-possible rate that might be expected from microevolutionary considerations (Lynch, 2011; Lynch et al., 2016). The finding is also significant because, while the mutational load has been considered to be a primary cause of genetic load (Klekowski, 1988), the possibility of a similarly sized impact for the latent load appears to have been rarely considered.

Despite the importance of the mutation rate to population mean fitness, and thus the success of species, there are limited microevolutionary mechanisms for keeping the mutation rate near its long-term population-optimal value. Consequently, if the mutation rate is to be kept close to its long-term population-optimal value, it will likely be necessary to invoke macroevolution and the extinction of species having mutation rates that are significantly too small or too large.

This manuscript builds on the work of Maynard Smith on the lag load (Maynard Smith, 1976), Haldane on the mutational load (Haldane, 1937), and Kimura et al. on the fixation of a beneficial allele, the genetic load in the presence of drift, and the heterozygosity of sites maintained by a flux of mutations (Kimura, 1962; Kimura et al., 1963; Kimura, 1969; Kimura and Ohta, 1969).

Kimura made the same argument as made in this manuscript that inter-group selection is responsible for the lower bound on mutation rates and he sought to determine the optimal mutation rate for the survival of a species, but the latent load was not considered (Kimura, 1960, 1967). It is argued in the present manuscript that the latent load is a key component in determining the population-optimal mutation rate.

It is beyond the scope of this manuscript to review the degree to which mutation rates are found to be close to their population-optimal values. This manuscript simply seeks to develop a mathematical model of population mean fitness, and then use it to derive a formula for the population-optimal mutation rate.

## Results

Let *w* denote the relative fitness of an organism’s genotype; that is its expected contribution to the next generation. Most of the time it is the relative nature of fitness that concerns us, but in order to speak meaningfully of fitness values alone it is necessary to anchor the fitness scale. Fitness here is defined relative to a hypothetical ideal organism that is perfectly adapted to its environment, and which is assigned a fitness value of 1. This means that log fitness is negative, or zero in the one special case. The concept of an ideal organism is a bookkeeping device rather than a realizable genotype.

As used here, genetic load describes the difference between the fitness of an ideal organism at a given point in time and the mean fitness of the current population. Traditionally,

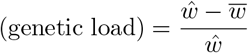

where 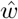 is the fitness of the ideal organism, which here has the value 1. The multiplicative nature of fitness makes this definition cumbersome to work with when combining multiple genetic loads unless they are all small. It is easier to consider the log of population mean fitness, log 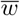, since logs combine additively.

Four contributors to the genetic load will be considered here:

- what will be termed the “latent load”, which results from sites that could harbor beneficial alleles but at which such alleles have not yet arisen and begun to fix;
- the substitutional load, which results when beneficial alleles have become established and the sites are in the process of fixing;
- the mutational load, which results from deleterious mutations;
- and the drift load, which results from genetic drift in the presence of mutations to nearly neutral sites in the genome.

The drift load could in principle be viewed as part of the mutational load. It is however convenient to treat it as a distinct load because unlike the mutational load it is independent of the mutation rate.

The environment is made up of both external and internal environments. The external environment is the aggregate of all the physical and biological spaces with which organisms making up the population interact. The internal environment is made up of the population’s genomes. The latent and substitutional loads are generated as a result of ongoing changes in the external environment. If the external environment were constant the latent and substitutional loads would tend toward zero. The rate of external environmental change may vary over time.

Many alleles are conditionally beneficial, conditionally deleterious, or conditionally neutral, exhibiting different beneficial, deleterious, or neutral effects depending on the allele’s environmental context. Such alleles are simply described here as beneficial, deleterious, or neutral depending on their aggregate effects across all organisms in the population. The effective selection coefficient of such an allele is the expected value of the allele’s selection coefficients across all organisms.

Figure 1 shows a simplified lifecycle of a site, and the different genetic loads it may generate. In this manuscript, a positively selected site is defined as a site for which there is an allele with a selection coefficient that is beneficial which has not previously been fixed, irrespective of whether the beneficial allele is currently present in the population. This definition is consistent with a conserved, or negatively selected site, not requiring deleterious alleles be present in the population. The definition differs from definitions of a positively selected site that require currently segregating beneficial alleles, or that simply require the site underwent positive selection at some point in the past.

**Figure 1:**
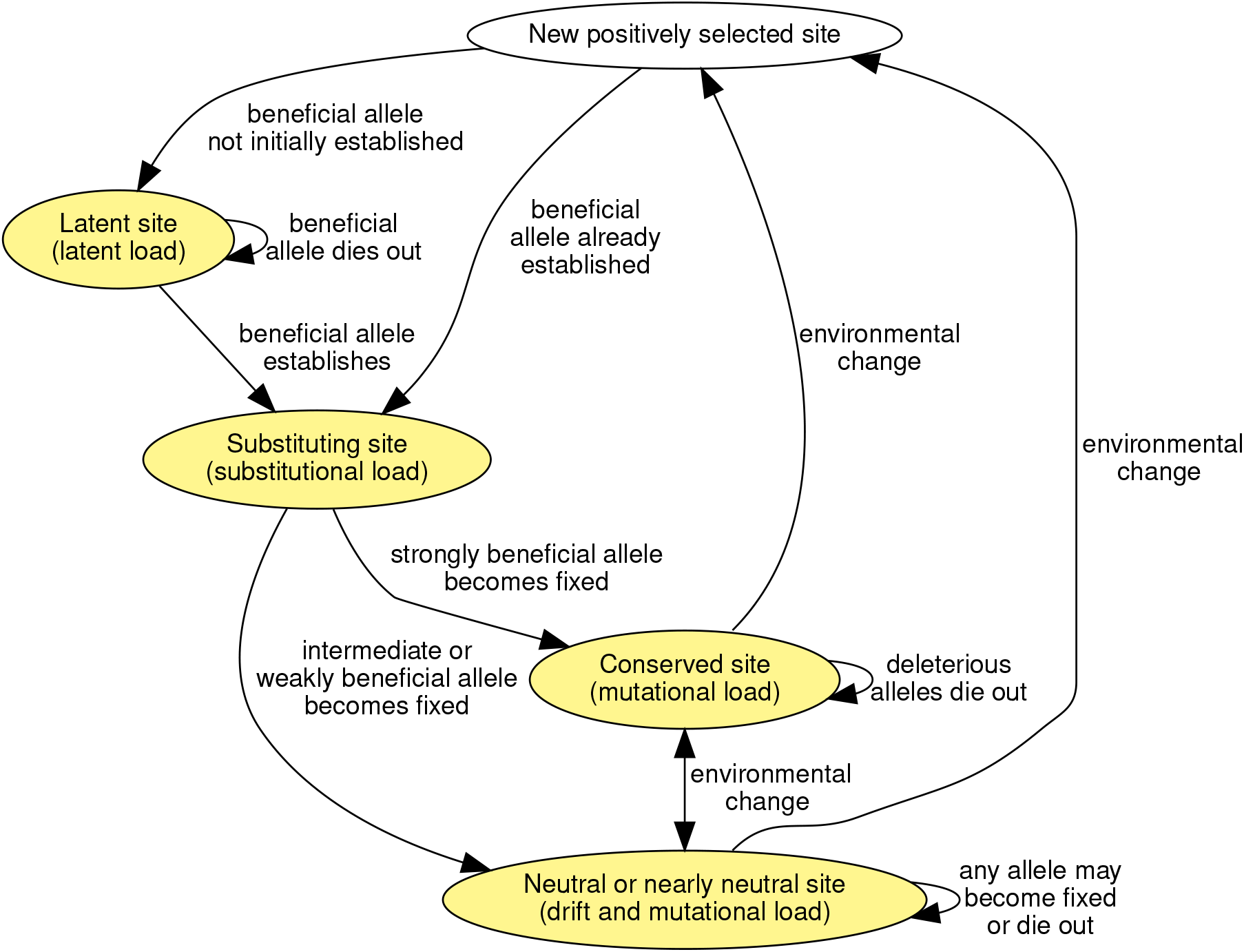
Simplified lifecycle of a site. Transitions between states are the result of mutation, selection, drift, and environmental change. New positively selected site is a transitory state.

A positively selected site is either a latent site or a substituting site. A beneficial allele is considered established if it is not going to die out, a fact that is only known with certainty in hindsight. A new positively selected site may have a beneficial allele that is already established as a result of standing genetic variation, or more typically, the beneficial allele first needs to arise by mutation and then become established. While waiting for the beneficial allele to establish the positively selected site is a latent site and it generates a latent load. Once the beneficial allele is established the positively selected site is a substituting site and it generates a substitutional load. When the beneficial allele becomes fixed the site is either conserved and generates a mutational load, or nearly neutral and generates a drift and mutational load. Occasionally a change in the environment causes a site to transition between being a conserved site, a neutral or nearly neutral site, or to become a new positively selected site.

The latent and substitutional loads together comprise what Maynard Smith calls the lag load (Maynard Smith, 1976). They exist because the makeup of the population lags behind that of an ideal organism as dictated by the current external environment, which changes over time. Maynard Smith’s concept of an ideal organism was as the locally optimal organism reached by a series of fitness increasing mutational steps should the external environment be held constant. The motivation for considering the latent and substitutional loads separately is that the latent load depends on the mutation rate, while the substitutional load is independent of the mutation rate.

### Model

A summary of the key notation used is provided in Table 1. Assume a sexual population of organisms with population size *N*, and apparent variance effective population size *N*_*e*_. Let the ploidy, *P*, be 1 for organisms that are counted and experience the force of selection during the haploid state, and 2 for organisms that are counted and experience selective forces while in the diploid state.

**Table 1:**
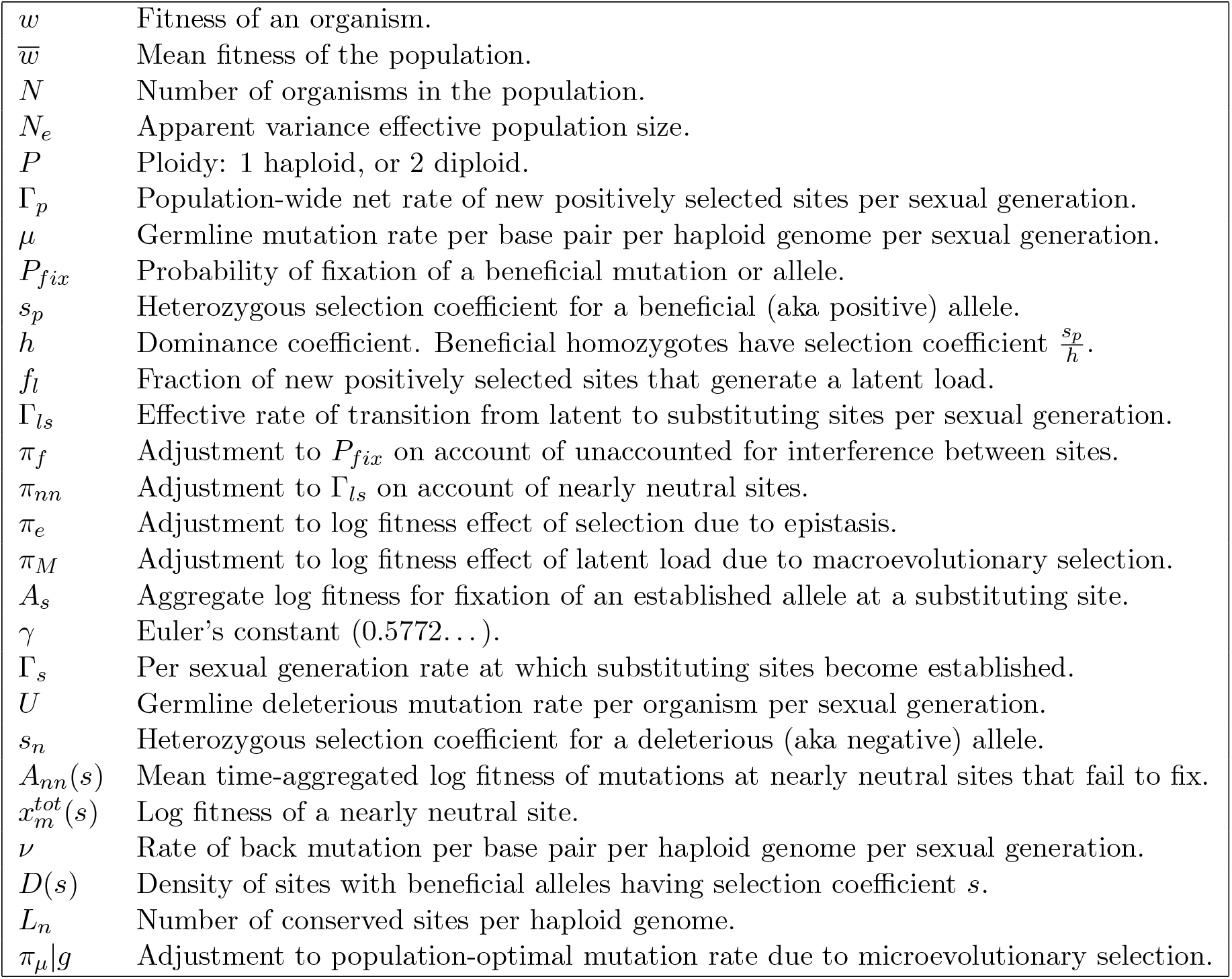
Summary of key notation.

Only single nucleotide mutations are considered. Insertion and deletion (indel) mutations are not included in the analysis.

Epistasis is possible, but epistasis among simultaneously segregating sites is assumed to be minimal. That is, a site that is either a latent site or a conserved site may affect the fitness of another site, but there should be few interactions between the fitness effects of substituting sites or nearly neutral sites.

Strong selection occurs when |*PN*_*e*_*s*| ≫ 1, where *s* is the selection coefficient. Sites with alleles having selection coefficients of intermediate strength behave according to the nearly neutral site model. In fact, the conserved site model is simply a special case of the more general nearly neutral site model.

In the model changes in selection coefficients occur as a result of environmental change, and these changes can be instantaneous or gradual. Latent sites are modeled as coming about as a result of instantaneous changes from neutral, nearly neutral, or conserved sites to sites with alleles having positive selection coefficients. If the new positive selection coefficient is strong this may be followed by further gradual variation in the strong positive selection coefficient of the latent site’s allele. The effects of gradual transitions by latent sites over the non-strong selection coefficient range are not modeled. Changes in the selection coefficients at substituting sites are not modeled. This is acceptable as the substitutional load is shown to be much smaller than the latent load.

The selection coefficient of a beneficial allele at a latent site may cease to be beneficial before the site becomes established. The effect of this is handled in the model by dealing with rates of transition from latent to substituting sites. This detail is not shown in Figure 1.

A site may have an associated selection coefficient so small that fixation of the beneficial allele would take longer than the lifetime of the species, but such sites are still relevant. This is because there are likely to be many such sites, and so it is possible to speak in terms of probabilities and rates of fixation.

### New positively selected sites

The external environment may undergo change either frequently or infrequently. When it does, such change may reduce the fitness of an organism relative to the ideal organism, which is assumed to have adapted to the change. Such change, however, creates opportunities for beneficial alleles to arise and/or increase in frequency at sites where the allele was previously neutral or deleterious. In addition to new positively selected sites arising as a result of changes in the external environment, new positively selected sites may also arise as a result of changes to the internal environment. The fixation of the beneficial allele at one positively selected site might epistatically lead to the creation of other nearby or distant positively selected sites.

Let Γ_*p*_ denote the net rate at which new positively selected sites arise per sexual generation.

Γ_*p*_ is a net rate because some positively selected sites may cease to be positively selected as a result of environmental change. It is also a net rate because a reduction in *N*_*e*_ may render some positively selected sites as nearly neutral, and thus reduce Γ_*p*_. Similarly, an increase in *N*_*e*_ may make some nearly neutral sites positively selected sites, increasing Γ_*p*_. By notational convention, a change in the selection coefficient of an already fixed allele to be beneficial will be excluded from Γ_*p*_. For example, if a site was neutral and fixed at allele B, and then the environment changes and allele B is now beneficial, this does not contribute to Γ_*p*_. Since base pairs can take on four possible values, this reduces the magnitude of Γ_*p*_ by roughly 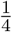.

### Probability of a latent site

Let *µ* denote the mean germline spontaneous mutation rate per base pair per haploid genome per sexual generation.

When a new positively selected site arises, its fitness impact will depend on whether the beneficial allele is already established and will proceed to fixation, or whether it either does not exist at all or existing copies of the allele will die out.

Recall that *P* denotes the ploidy. Let *s*_*p*_ denote the heterozygous selection coefficient associated with a beneficial allele. Based on the work of Kimura (Kimura, 1962; Patwa and Wahl, 2008), if the allele has proportion *p*, the probability of fixation, *P*_*fix*_, is,

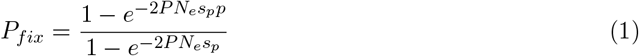

Thus if 2*PN*_*e*_*s*_*p*_*p* ≫ 1 the beneficial allele at the previously conserved or neutral site is already established, and so the site will only ever exert a substitutional load. Conversely, if 2*PN*_*e*_*s*_*p*_*p* ≪ 1, the probability of fixation of the beneficial allele is small, and so the substitutional load is preceded by a latent load.

Let *f*_*l*_ denote the probability that a new positively selected site is a latent site. That is, at the time the new positively selected site arises all existing copies of the beneficial allele are destined for extinction. Appendix A shows that *f*_*l*_ will have a value close to 1 (e.g., *f*_*l*_ = 0.992 for *PN*_*e*_ = 2 *×* 10^5^, *µ* = 10^−8^, and *s*_*p*_ = 10^−3^ which are values typical of vertebrates).

### Latent load

The genetic load associated with a new positively selected site does not begin at the time that a beneficial allele establishes, but at the time the environment changes to make an allele’s selection coefficient positive, thereby creating a new positively selected site. The genetic load from the time a new positively selected site comes into existence until the beneficial allele is established is termed the latent load.

For simplicity, it is assumed that only one of the four possible base pair values will maximize the fitness associated with a new positively selected site, and that the other base pair values are equally deleterious. Consequently, the mean rate of beneficial mutations per haploid genome at a latent site is approximately 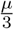.

For now, assume that there is no epistasis.

Also for now, assume that environmental change causes new positively selected sites to arise in an all-or-nothing manner; a site is initially neutral, nearly neutral, or conserved, and then suddenly becomes a positively selected site with a fixed selection coefficient *s*_*p*_ for the beneficial allele.

Figure 2 shows the rates of transition between different types of positively selected sites.

**Figure 2:**
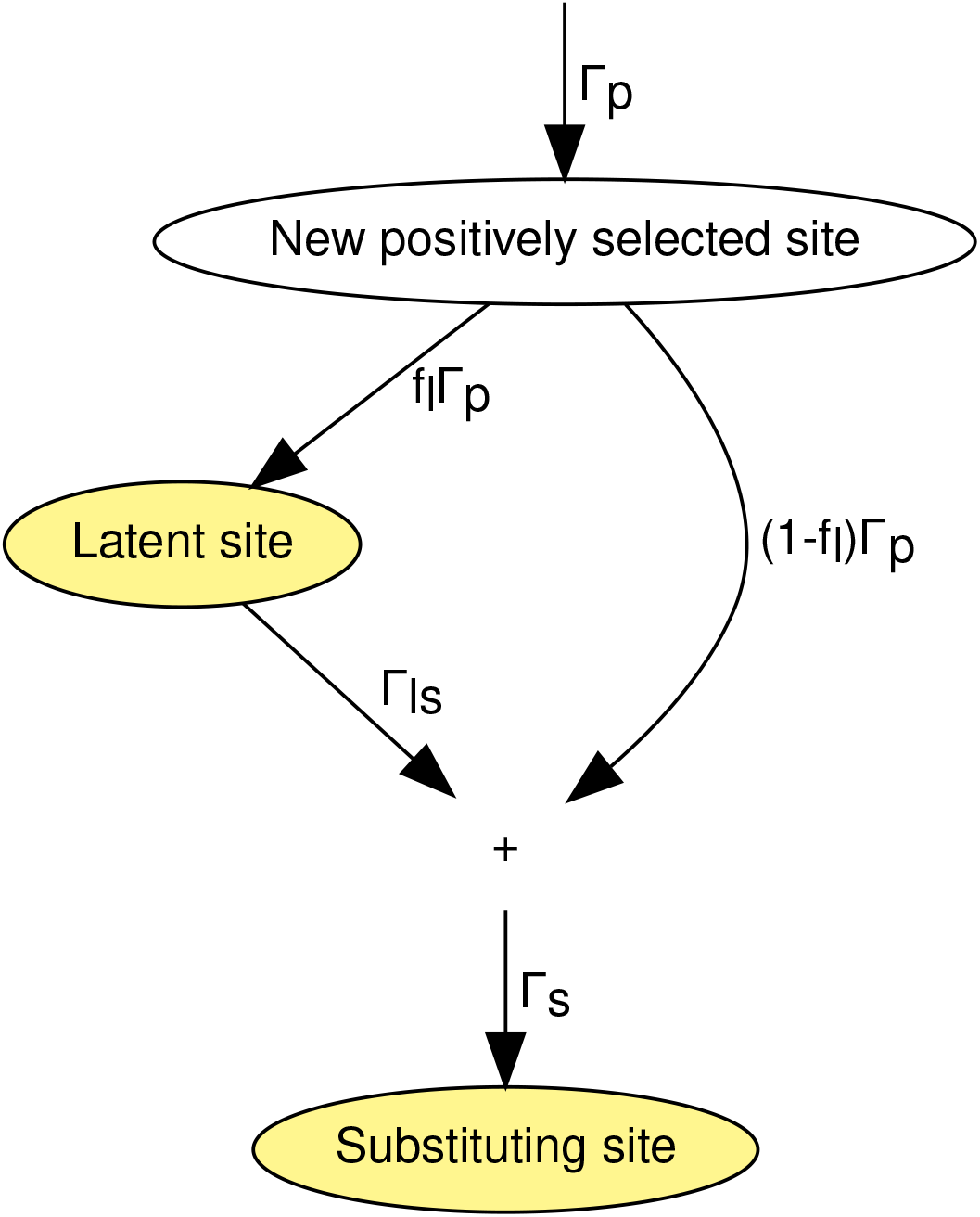
Rates of transition between different types of positively selected sites.

Let Γ_*ls*_ denote the rate at which latent positively selected sites transition to being substituting sites.

Let Γ_*s*_ denote the per sexual generation rate at which substituting sites become established. Substituting sites become established either when a beneficial mutation of a latent site establishes, or when the beneficial allele of a new positively selected site is already established. Thus,

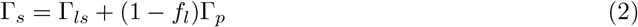

For a species pair, over macroevolutionary timescales, the temporal mean value of Γ_*s*_ is equal to the rate of adaptive substitution per sexual generation, and can be estimated by,

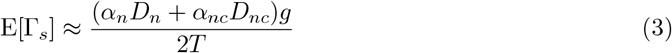

where *α*_*n*_ is the fraction of non-synonymous substitutions that are fixed by positive selection, *D*_*n*_ is the number of non-synonymous substitutions, *α*_*nc*_ is the fraction of substitutions that are fixed by positive selection in the non-coding region, *D*_*nc*_ is the number of substitutions in the non-coding region, *g* is the generation length, and *T* is the divergence time. The fixation by drift of sites with alleles having small fitness effect is corrected for the determination of *α*_*n*_ and *α*_*nc*_.

Suppose that genome-wide there are a total of *k*_*l*_ latent sites. Each site exists in every haploid genome. Latent sites arise at a rate of *f*_*l*_Γ_*p*_. The difference between these two rates is equal to the rate at which *k*_*l*_ changes per sexual generation, Δ*k*_*l*_,

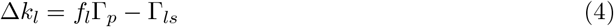

Eliminate Γ_*p*_ by combining equations 2 and 4,

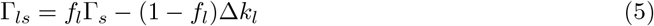

Fixation of the beneficial allele at a latent site typically starts from a single mutation. Thus, in this case, when *s*_*p*_ ≪ 1, from equation 1,

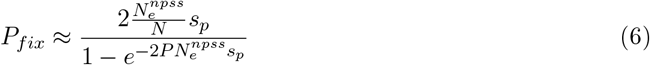

where 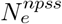 is the effective population size at the new positively selected site if it were instead neutral.

The aggregate rate of beneficial mutation at the latent sites is *k*_*l*_*PN* 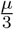, and the probability of fixation of each beneficial mutation is approximately given by equation 6. The product of these two terms equals the rate at which latent sites transition to substituting sites,

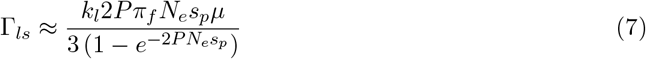

where *π*_*f*_ is a factor to account for any change to the probability of fixation on account of a possible difference between the average effective population size of new positively selected sites, 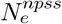, and the average effective population size across all sites, *N*_*e*_. If, as is likely, new positively selected sites cluster in time and space the two may differ; differences in the expected effects of linkage will then need to be taken into account. Appendix B shows that *π*_*f*_ is expected to be close to 1 (e.g., *>* 0.99 for apes and rodents, and *>* 0.97 for insects).

Let *h* denote the dominance coefficient. By convention *h* = 1 for haploid eukaryotic species. Assuming *s*_*p*_ ≪ 1, the log fitness loss associated with a wild-type homozygote is approximately 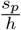. This is the log fitness gained if the organism is a homozygous mutant. The wild-type homozygote contributes to the latent genetic load.

Let log 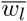 denote the log of the mean fitness associated with latent sites,

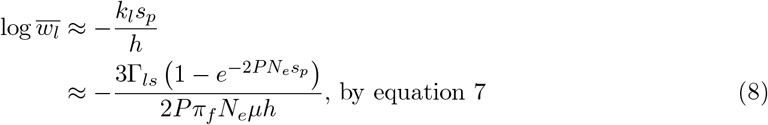

In the event that *h* is not constant but instead a function of the latent sites, *h* in equation 8 represents the harmonic mean value.

If *s*_*p*_ is strong, the exponential term is approximately zero, and equation 8 is effectively independent of *s*_*p*_ and equally applicable to a distribution of *s*_*p*_ values. When *s*_*p*_ is nearly neutral the exponent term will be close to 1, the log fitness effect will be proportional to *s*_*p*_, and it will be much smaller than usual. Let Γ_*ls*_(*s*) denote the density of the rate of transition from latent to substituting sites with beneficial alleles having selection coefficient *s*. For a range of selection coefficients define the latent to substituting site nearly neutral adjustment *π*_*nn*_ by,

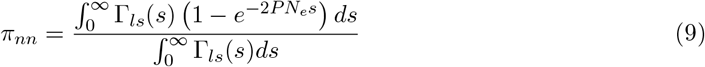

Then,

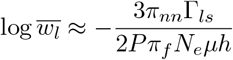

It is shown in Appendix C that for three animal species that were examined, assuming an exponential distribution of fitness effects at latent sites, 0.96 *< π*_*nn*_ ≤ 1.

Epistasis is likely to significantly alter the number of current positively selected sites that actually fix. There are many genotypic paths to the same phenotype. By redefining the latent load to be the fitness effect only of positively selected sites that are destined to fix, Appendix D shows how it is possible to incorporate epistasis into the latent load. To reflect epistasis, let,

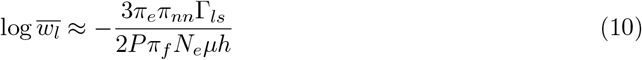

where *π*_*e*_ *>* 0 is an adjustment to the relative log fitness effect of selection on account of epistasis. Appendix D shows an estimate of *π*_*e*_ is 0.8; approximate range 0.6 to 0.9.

Since *f*_*l*_ is close to 1, barring Γ_*p*_ becoming orders of magnitude larger than Γ_*s*_, which is not realistic, (1 − *f*_*l*_)Δ*k*_*l*_ ≈ 0. Thus by equation 5,

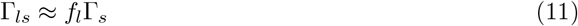

Equation 10 is valid for a population at a given point in time. Also of interest is the mean value of log 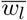 over an evolutionary time period, 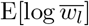. If the generation length is variable then 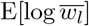 would be taken by averaging over the number of generations.

Equation 10 is true from a microevolutionary perspective, but not necessarily from a macroevolutionary perspective. Over time the number of latent sites for a population slowly fluctuates as new latent sites are created and existing sites transition to substituting sites. If the population is one of many that are in competition, then species with temporarily larger-than-usual numbers of latent sites will be less fit than usual, and thus more vulnerable to macroevolutionary extinction at that point. This would introduce a downward bias in the number of latent sites in extant species. To reflect this, define 0 *< π*_*M*_ ≤ 1, as an adjustment to the log fitness effect of the latent load on account of any macroevolutionary selection. Then,

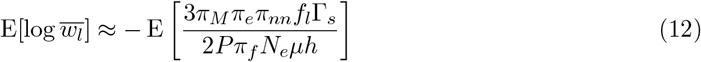

Thus far, the only case that has been considered, is one for which alleles at sites that become new positively selected sites instantly go from having a neutral or deleterious effect, to having a fixed positive selection coefficient. It is likely that, more commonly, the selection coefficient changes slowly over time as the environment changes, until it reaches its maximal value. Appendix E demonstrates that provided the allele is strongly beneficial, this gradualism is unlikely to alter the value of log 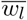

### Substitutional load

It takes time for the beneficial allele at a positively selected site to reach fixation. From the moment a beneficial allele is established until it is fixed, it exerts a substitutional load.

Appendix F derives the following approximation for the fitness effect of the substitutional load, log 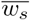, in the presence of strong selection,

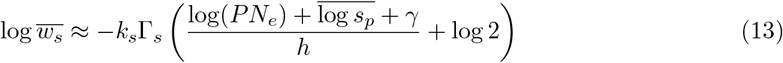

where *k*_*s*_ denotes a value less than or equal to 1 that reflects preexisting established beneficial alleles of new positively selected sites, and *γ* is Euler’s constant (0.5772…).

### Mutational load

Mutations at conserved sites are deleterious and are removed from the population either by selection or by chance back mutation. Suppose deleterious mutations occur randomly with a mean germline rate *U* per organism per sexual generation. Let −*s*_*n*_ denote the heterozygous selection coefficient associated with deleterious alleles, with *s*_*n*_ *>* 0. If *s*_*n*_ ≫ *µ*, and 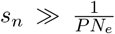, so that selection dominates over drift, then Haldane has shown (Haldane, 1937; Agrawal and Whitlock, 2012),

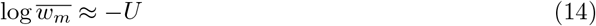

### Numerical comparison of loads

For humans, it appears that in log terms the latent load is larger than the mutational load, and more than two orders of magnitude larger than the substitutional load.

An estimate of Γ_*s*_ for humans and chimps can be obtained by using equation 3 with *α*_*n*_ = 0.10 − −0.13 (Gojobori *et al*., 2007), *D*_*n*_ = 3.9*×*10^4^ (Sequencing and Consortium, 2005), *α*_*nc*_*D*_*nc*_ = 0 (Keightley *et al*., 2005), *g* = 26 years (Wang *et al*., 2023; Langergraber *et al*., 2012), and *T* = 6.4*×*10^6^ years (Kumar *et al*., 2017), to give E[Γ_*s*_] ≈ 9.1*×*10^−3^. For humans *µ* ≈ 1.2*×*10^−8^ and the neutral site *N*_*e*_ ≈ 2.2*×*10^4^ (Lynch *et al*., 2023)[EV1].

For the latent load, using the above values in equation 12 along with *π*_*M*_ = 1, *π*_*e*_ = 0.8, *π*_*nn*_ = 1, *f*_*l*_ = 1, *P* = 2, *π*_*f*_ = 1, and *h* = 0.8 gives 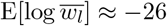.

For the substitutional load, using the same values in equation 13 along with *k*_*s*_ = 1 and *s*_*p*_ = 10^−2^ gives log 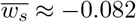.

For the mutational load, the median estimate of the fraction of the genome under the control of negative selection, *f*_*n*_, has been reported as 5.5% (Ponting and Hardison, 2011). Using equation 14 and *U* = *Pf*_*n*_*Lµ* with the genome length, *L* = 3.1*×*10^9^, gives log 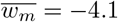.

Because the latent load and the substitutional load both involve the same factor Γ_*s*_, and because *N*_*e*_*µ* ≪ 1 for nearly all sexual eukaryotes (Lynch *et al*., 2023)[EV1], the latent load is expected to be far larger than the substitutional load for nearly all sexual eukaryotes.

### The population-optimal mutation rate

The likelihood of a mutation being harmful is far greater than the likelihood that it is beneficial. Further, the magnitude of the fitness effect of harmful mutations is typically greater than that of beneficial mutations. Consequently, the determinant of the lower bound on the mutation rate in sexual organisms has been something of a mystery (Leigh Jr, 1970; Johnson, 1999). One hypothesis is that natural selection drives the mutation rate down to a lower bound set by the balance between selection for lower mutation rates and random genetic drift (Lynch, 2011; Lynch et al., 2016).

Here it is proposed that, in the long run, evolution biases the germline spontaneous mutation rate per sexual generation so that it is as small as possible to minimize the mutational load, but not so small that the latent load becomes excessively large. That is, evolution results in a mutation rate that approaches a species or population-optimal value. This is distinct from the optimal mutation rate for an individual organism or its genes. This process may occur through macroevolution. Species that evolve mutation rates that are too low fail to adapt, incur a larger latent load, and are more likely to become extinct. Species with mutation rates that are too high incur a heavy mutational load, produce fewer successful offspring, and are likewise more likely to become extinct.

At first glance, it may seem implausible that species go extinct due to suboptimal mutation rates. But this is proposed to be only a contributing factor. Other stochastic factors also contribute to the success of a species. But in the same way that the selective advantage of a new genotype does not determine the realized number of copies in the next generation, but only the odds of copies, the optimality of the mutation rate affects the odds of species survival. Over macroevolutionary time scales these odds can be expected to accumulate, leading to mutation rates in the surviving species that are roughly appropriate for the relevant lineages.

An analogy might be drawn between the proposed macroevolution of sexual species with variable mutation rates, and the observed microevolution of asexual organisms with variable mutation rates. A sexual species is viewed as analogous to a single asexual organism. In bacteria when there is a significant selective challenge, and a higher mutation rate is desirable, mutator alleles that increase the mutation rate frequently arise and occasionally spread by hitchhiking with beneficial mutations (Sniegowski *et al*., 1997). Once the challenge has been met, the mutation rates subsequently decline (Wielgoss *et al*., 2013). For this analogy to be valid there needs to be some mechanism for species to occasionally acquire higher mutation rates simultaneously with increased population mean fitness. Whether this is possible through stochastic effects in small species founder populations is not certain. Another mechanism through which a higher mutation rate might be possible is if the species evolves a longer lifespan. Other things being equal, a longer lifespan might frequently be expected to increase the per generation mutation rate, while at the same time increasing the number of offspring and thus offering a possible fitness boost. This mechanism also is not certain as there are complications in so far as a longer lifespan might also reduce the population mean fitness of the progeny.

Let *L*_*n*_ denote the number of sites under the control of negative selection per haploid genome, so that,

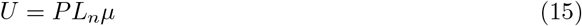

Strictly speaking, in what follows, *L*_*n*_ and *U* should be considered effective quantities, reflecting not just conserved sites but also the partial contribution to the mutational load of nearly neutral sites. The genetic load of nearly neutral sites is discussed in Appendix G.

Let log 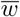 denote the log of mean fitness of a population. The only genetic loads which depend on the mutation rate are the latent load and the mutational load. As shown in Appendix G, the drift load is only dependent on the ratio of forward to back mutation. Combining equations 10 and 14,

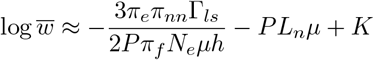

where *K* consists of terms that are independent of *µ*.

When combining multiple site types, it is important to consider possible interference between their fitness effects. Interference occurs when the presence of one site affects the fitness effect of another. There is no interference from the latent load on the mutational load because, apart from rare copies of beneficial alleles that die out, there is no variation at the latent site. Interference by the mutational load on the latent load is captured by the mutational load’s effects on the effective population size.

Suppose *P, L*_*n*_, and *µ* are fixed, but that the other parameters are functions of time.

A population is most likely to be successful if the product of successive 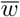 terms is as large as possible, or equivalently the sum of their logs is maximized. This is the same as maximizing 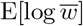. Using equation 12,

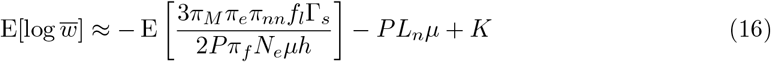

Differentiating equation 16 with respect to *µ* shows that the geometric-mean population mean fitness is maximized when,

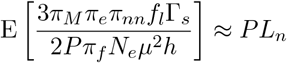

So the maximal population fitness is given by 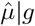, where,

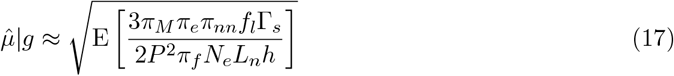

The given *g* qualifier signifies this value is for a given generation length. It is not necessarily the optimal value if the generation length, and thus Γ_*p*_ and Γ_*s*_, are free to also change.

Equation 17 gives the fixed mutation rate that would be near optimal for the species over the time frame (measured in generations) for which the expectation was taken. Care must be taken in applying equation 17 if the parameter values vary over time; it might not be possible to accurately drop the expectation operator.

It should be obvious that species do not know in advance the parameter values for the optimal mutation rate; rather the equation for population mean fitness acts as a mutation rate “filter” through which competing species must pass.

The maximal log fitness is given by,

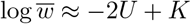

At the population-optimal mutation rate, mean fitness losses from the latent load and mutational load are approximately equal.

Microevolution might drive *µ* to a value below than the population-optimal value. Let *π*_*µ*_|*g* denote the multiplicative factor by which the mutation rate is smaller or larger than its optimal value. Then,

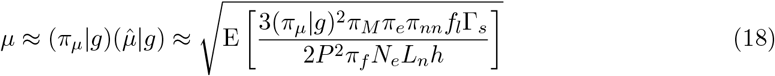

and,

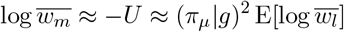

Most of the terms in the right hand side of equation 18 are expected to have similar values within very broad clades (Linnaean kingdoms). The exceptions are *L*_*n*_, which might show some *N*_*e*_ dependence but should still be largely constant within slightly less broad clades (Linnaean phyla), *N*_*e*_, and lastly Γ_*s*_, which might like Γ_*p*_ exhibit considerable stochastic variation.

Ignoring the temporal effects of the expectation operator, equation 18 suggests an inverse (square root) relationship between population sizes and mutation rates for sexual species. Species with large population sizes such as invertebrates, and especially unicellular eukaryotes, are expected to have smaller mutation rates per sexual generation than vertebrates. This appears to agree with empirical observations (Lynch *et al*., 2023). Assuming other parameters are held constant, equation 18 predicts the log mutation rate equals 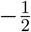 times the log effective population size, plus a constant, plus noise from the variation in Γ_*s*_. This also approximately agrees with empirical observations both within and between clades (Lynch *et al*., 2016, 2023).

## Discussion

A new load, the latent load, was isolated from the lag load and analyzed. The latent and mutational loads imply the existence of a spontaneous mutation rate per base pair that maximizes the aggregate mean fitness of a population. If the mutation rate is too low, beneficial alleles take a long time to establish, increasing the latent load. If it is too high, negative selection incurs a large mutational load. The population-optimal mutation rate is one at which the mean fitness losses from the latent and mutational loads are approximately equal. The extent to which evolution approaches the population-optimal mutation rate is a question for further study.

## Appendix A: The latent fraction of new positively selected sites

The latent load depends on the value of *f*_*l*_, the proportion of new positively selected sites for which no existing copies of the beneficial allele are already established.

For a new positively selected site that arises at a previously neutral site, *f*_*l*_ can be estimated as follows.

Assume the beneficial allele suddenly has selection coefficient *s*_*p*_, and all other possible alleles have a selection coefficient of zero.

From equation 1, assuming 2*PN*_*e*_*s*_*p*_ ≫ 1, the probability of fixation of an allele present at proportion *x* is given by,

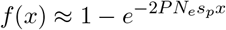

The work of Kimura provides a method for determining the mean value of *f* (*x*) (Kimura, 1969). Adopting Kimura’s notation, let,

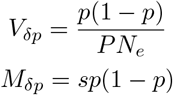

for some heterozygous selection coefficient *s*, with dominance coefficient, 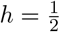. In a moment, the limit as *s* → 0 to describe a previously neutral site will be considered. Let,

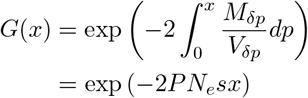

Let,

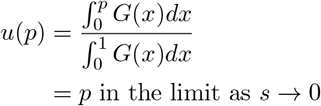

If the mutation rate is *µ*, and one third of mutations produce the currently neutral, but soon-to-be beneficial allele, and *ψ*_*f*_ (*ξ*) is defined by,

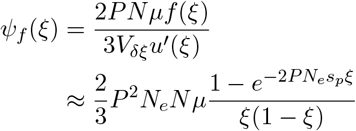

and *I*_*f*_ (*p*) is defined by,

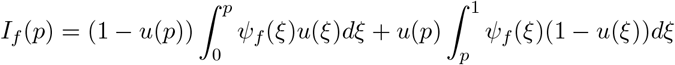

then

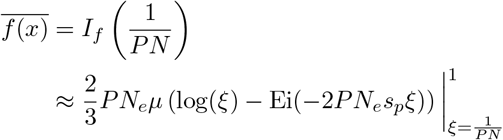

because the first term of *I*_*f*_ 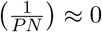 when *PN* ≫ 1, where Ei is the exponential integral,

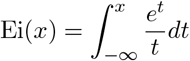

Now, Ei(*x*) ≈ 0 for *x* ≪ −1. In addition, Ei(*x*) ≈ *γ* + log(|*x*|) for |*x*| ≪ 1 where *γ* is Euler’s constant (0.5772…). Thus, assuming 2*PN*_*e*_*s*_*p*_ ≫ 1 and 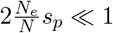,

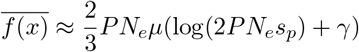

This is the probability of fixation of the now-beneficial allele at a site that was neutral and suddenly became a new positively selected site. The latent fraction, *f*_*l*_, is 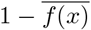, or,

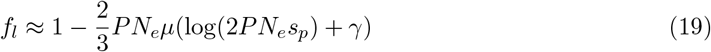

Now consider a distribution of selection coefficients. Suppose an exponential distribution of fitness effects for new positively selected sites with mean selection coefficient 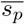. The exponential distribution is chosen here for analytical convenience. From equation 9 the latent load is likely to be substantial when 2*PN*_*e*_*s*_*p*_ ≳ 1. Evaluating the latent fraction for just those sites,

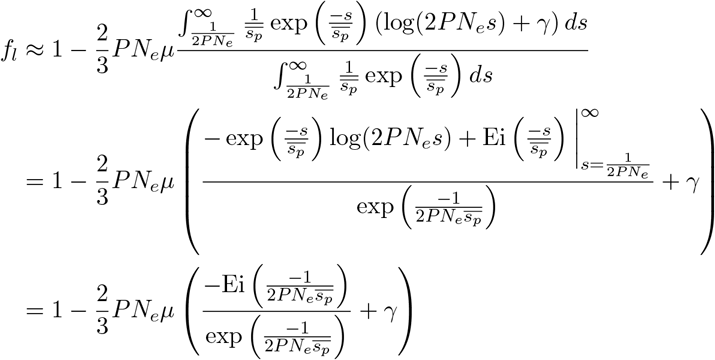

For values typical of vertebrates (*PN*_*e*_ = 2*×*10^5^, *µ* = 10^−8^, and 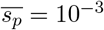), this yields *f*_*l*_ = 0.992.

This analysis assumes 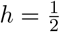; other dominance coefficients may change the scaling but are unlikely to alter the qualitative result.

This result applies to a new positively selected site that arises at a previously neutral site. If the new positively selected site arises at a site where the beneficial allele was previously deleterious, the initially expected proportion of the beneficial allele is smaller, making *f*_*l*_ even closer to 1.

## Appendix B: Clustering of new positively selected sites

Adaptation to an environmental change frequently appears to consist of an adaptive walk of three to six discrete steps (Schoustra *et al*., 2009), rather than a long series of steps where each step is triggered by the preceding step. It does not appear to be known whether the ordering of these steps is largely arbitrary, or whether the ordering is fixed due to sign epistasis between adaptive sites. It also does not appear to be known whether the adaptive sites are physically located close to each other on the genome. If the ordering of the steps is arbitrary and the sites are physically close, then a beneficial allele undergoing a selective sweep at one site may interfere with and reduce the probability of fixation of a beneficial allele at another.

Most interference between sites is captured by *N*_*e*_. However, the effect of the clustering of new positively selected sites will not be captured by *N*_*e*_ because *N*_*e*_ is the effective population size averaged across the entire genome, rather than the local effective population size experienced by tightly linked positively selected sites. This reduction in the probability of fixation is denoted *π*_*f*_. Since *π*_*f*_ ≤ 1, equation 12 evaluated without *π*_*f*_ may underestimate the mean latent load.

There is some concern that undetected new positively selected sites of smaller fitness effect than those detected in the studies of adaptive walks cited above might increase the mean length of an adaptive walk. However, as will be shown, if they are of smaller fitness effect, they are unlikely to overlap in time with the detected sites. Conceivably, such sites could then overlap in time with each other, reducing their probability of fixation. This effect is not modeled here, but could contribute to a reduction in the estimate for *π*_*f*_.

The rate of adaptive amino acid substitution at a given location in the genome has been reported to be positively correlated with the mutation rate and the recombination rate, and negatively correlated with gene density in *Drosophila* (Castellano *et al*., 2016). Based on this it was estimated that genome-wide Hill-Robertson interference is responsible for the loss of about 27% of all adaptive substitutions. However, the interpretation of this result is not straightforward within the model used in this manuscript. If a beneficial mutation fails to establish and ultimately fix, then the latent site will remain, beneficial mutations will reoccur, and ultimately establish and fix. In the model used here, adaptive substitutions are delayed rather than permanently lost, because failure to establish leaves the site as a latent site rather than eliminating future adaptive opportunity.

Barton has calculated the probability of fixation of a strongly selected beneficial mutation at a site that is tightly linked to another site that is undergoing a selective sweep (Barton, 1995). For a beneficial mutation with an effect size much smaller than that of the sweeping allele, and occurring prior to the sweeping allele reaching a frequency of 50%, the reduction in fixation probability can be substantial.

Consider the probability that two potential selective sweeps at two nearby new positively selected sites overlap critically in time.

Suppose there are two sites: a first site harboring a beneficial allele with selection coefficient *s*_*p*_, for which *PN*_*e*_*s*_*p*_ ≫ 1, and a second site with a beneficial allele with selection coefficient *S*_*p*_, *PN*_*e*_*S*_*p*_ ≫ 1 that is destined to undergo a selective sweep. Assume the two sites are linked. Also assume the two sites were created as a result of the same environmental change, and that initially both sites have a frequency of zero for their respective beneficial alleles. Assume the second site establishes first; otherwise, the roles of the two sites may be interchanged.

Suppose 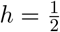. The time from the population acquiring a single copy of the second beneficial allele until it reaches the critical value of 50%, *T*_*S*_, is approximately given by (Otto and Whitlock, 2005),

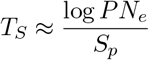

Let *T*_*l*_ denote the time, measured from the establishment of the second site, until the first site would have become established in the absence of the second site. Beneficial mutations at the first site are occurring at rate 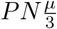, and the probability of fixation is normally 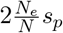. Consequently *T*_*l*_ has an exponential distribution with rate parameter 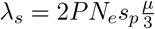.

Let *P*_*crit*_ denote the probability that the first site would have become established after the second site, but prior to the second site reaching a frequency of 50%; this is given by,

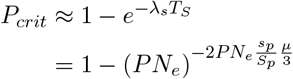

For *s*_*p*_ = *S*_*p*_ and *P* = 2, this gives *P*_*crit*_ ≈ 0.28% when *N*_*e*_ = 2*×*10^4^ and *µ* = 1*×*10^−8^, *P*_*crit*_ ≈ 1.7% when *N*_*e*_ = 2*×*10^5^ and *µ* = 5*×*10^−9^, and *P*_*crit*_ ≈ 9.2% when *N*_*e*_ = 1*×*10^6^, and *µ* = 5*×*10^−9^, values intended to be roughly representative of apes, rodents, and insects, respectively. These values are likely overestimates, because the fact that the second site established first typically means *s*_*p*_ *< S*_*p*_.

*P*_*crit*_ gives the probability of a potentially critical interaction between the two sites. This potentially critical interaction only applies to the first site. The second site is already established, and certain to fix. Selecting one of the two sites at random, the probability of a potentially critical interaction is 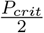. The extent to which this then reduces the probability of fixation of the first site will depend on the ratio of the selection coefficients, the degree of linkage between the sites, and the time at which the beneficial mutation which would have been establishing occurs. Consequently, the mean reduction in the probability of fixation, consequent reduction in the effective population size, and thus increase in the latent load, might be at most a few percent.

Thus, tight linkage between two positively selected sites will only infrequently affect their probability of fixation. The mean number of sites actually interacting is limited by adaptive walks that typically involve three to six steps, not all of which may potentially occur concurrently, and not all of which are at sites located close to each other on the genome. Hence, clustering of new positively selected sites appears unlikely to appreciably increase the latent load, and *π*_*f*_ is expected to be close to 1 (e.g., *>* 0.99 for apes and rodents, and *>* 0.97 for insects).

The population-optimal mutation rate is obtained by taking the square root of an expression involving the effective population size; equation 17. Thus, the population-optimal mutation rate will be increased by clustering of new positively selected sites by an even smaller proportion than the latent load.

## Appendix C: Adjustment to the latent load on account of nearly neutral sites

Let *ρ*_*p*_(*s*) denote the distribution of fitness effects for the beneficial alleles of sites transitioning from latent to substituting sites with selection coefficient *s*. By equation 11 this distribution is approximately the same as the distribution of fitness effects for new substituting sites. Theory suggests *ρ*_*p*_(*s*) may be modeled as an exponential distribution with mean fitness effect size 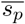 (Orr, 2003),

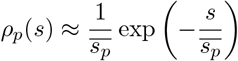

Note that *ρ*_*p*_(*s*) is not the same as the distribution of fitness effects for beneficial mutations that occur. Beneficial mutations of smaller effect size are less likely to fix; consequently, sites harboring them will persist as positively selected sites for longer, allowing the same beneficial mutation to recur more often prior to fixation.

Γ_*ls*_(*s*) is the distribution of fitness effects for sites transitioning from latent to substituting sites per sexual generation.

Let,

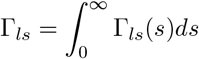

Then by definition,

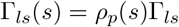

The nearly neutral adjustment, *π*_*nn*_, is defined by equation 9 as,

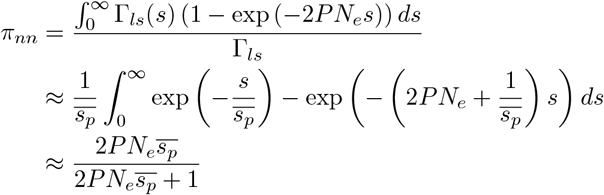

For humans, it has been estimated that 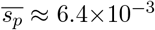, with a 95% confidence interval of 7*×*10^−4^ to 1*×*10^−1^ (Castellano *et al*., 2019), and *N*_*e*_ is of the order of 10,000. The distribution of fitness effects for positive selection in *Mus musculus* and *Drosophila melanogaster* has previously been modeled using a single fitness class with *PN*_*e*_*s*_*p*_ equal to 15 and 23, respectively (Booker, 2020). Consequently, *π*_*nn*_ equals 99.6%, 96.8%, and 97.9% for humans, mice and *D. melanogaster*, respectively.

## Appendix D: Epistasis and the latent load

Epistasis occurs when fixation of one positively selected site changes the selection coefficients of alleles at other sites. Several possibilities arise. An affected site may be a latent site, in which case the selection coefficient of the beneficial allele may increase (positive epistasis), decrease (negative epistasis), or it even become negative (a form of sign epistasis). Alternatively, an affected site may be a conserved site. A change in the selection coefficient of an allele at a conserved site will normally not have any appreciable effect on population mean fitness. The one exception is sign epistasis, in which the selection coefficient of a previously deleterious allele becomes positive, thereby creating the potential for future adaptive change.

Epistasis involving substituting sites and nearly neutral sites is not modeled here. Epistasis between latent sites, and sign epistasis at conserved sites, is considered here.

For sign epistasis to occur, both the direction and magnitude of the change in the selection coefficient must be appropriate. Positive or negative epistasis require neither a specific direction nor a specific magnitude. It therefore seems likely that the effect of sign epistasis on the latent load will be of second order relative to that of positive and negative epistasis.

In the presence of positive or negative epistasis, the latent load is defined in terms of the change in fitness that would result from the fixation of all current positively selected sites that are destined to fix, were the environment held constant. Sites subject to final sign epistasis are excluded from this definition. It is impossible to hold the environment constant, evolution may not be deterministic, and it is only known retrospectively whether a positively selected site is destined to fix or instead to undergo sign epistasis and cease being a positively selected site. This may cause empirical difficulties in precisely determining the latent load, but it introduces no conceptual difficulties.

Let *π*_*e*_ denote the mean multiplicative adjustment to the log fitness effect of the beneficial alleles of latent sites that are destined to fix, so as to incorporate the cumulative fitness effects of positive and negative epistasis experienced by all the other latent sites that are destined to fix. It follows that *π*_*e*_ is also the multiplicative adjustment to the log fitness effect of the latent load that accounts for epistasis.

Mean negative epistasis between beneficial mutations appears to be common (Rokyta *et al*., 2011; Schenk *et al*., 2013; Chou et al., 2011; Wei and Zhang, 2019; Schoustra et al., 2016). Using colony size as a proxy for fitness, Schoustra et al. investigated the epistatic effects of single and double mutants for the loss of an unused, costly cellular function (Schoustra *et al*., 2016). For mutations of similar effect size, they found double mutants had a mean fitness epistatic effect size relative to the mean of the two single mutants fitness effects of -0.18 in a rich physical environment and -0.66 in a poor physical environment when likely cases of sign epistasis are removed (Schoustra *et al*., 2016)[Calculated from data supplement]. This is suggestive of a mean epistatic effect size, relative to the sum of the two log fitness effects of approximately -0.2.

At any point in time there might be many latent sites. However, not all are destined to fix; some undergo reciprocal sign epistasis in response to those that fix. As discussed in Appendix B, adaptive walks in response to an environmental change typically consist of three to six steps. Not all of these steps may be capable of occurring concurrently; some may only arise in response to others. Thus, latent sites can probably be arranged into very small groups corresponding to concurrently feasible sites of adaptive walks for particular phenotypic effects, with relatively little epistasis occurring between groups. If the mean size of these groups is approximately two, then based on the data of Schoustra et al., a reasonable estimate for *π*_*e*_ is 0.8, with an approximate range of 0.6 to 0.9.

## Appendix E: The emergence of positively selected sites and the latent load

Until this point, the model has assumed an allele is either neutral or deleterious, and then instantaneously becomes beneficial with a constant selection coefficient. The real world is unlikely to operate in this manner. More likely, the beneficial effect of the allele increases gradually over time.

Let the selection coefficient, *s*(*i*), of the beneficial allele at the site be a function of the number of generations, *i*, since the allele first began to confer a beneficial effect. Suppose selection is strong, *PN*_*e*_*s*(*i*) ≫ 1 for all *i*, so that the probability of fixation of a single copy of the allele arising at generation *j* is 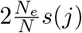.

The expected log fitness associated with the latent load of a single latent site of age *i* is given by the latent load for one period, multiplied by the product of the probabilities that the beneficial allele was not destined to fix in any prior period,

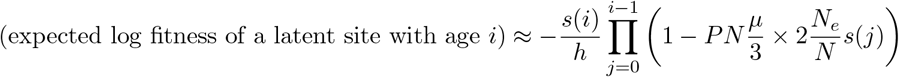

Then, the log of mean fitness associated with latent sites, log 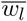, is given by,

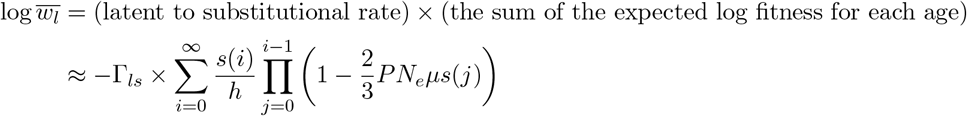

As expected, when *s*(*i*) = *s*_*p*_ for all *i*, this yields,

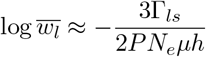

Let *r* denote the ratio of log 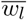for an arbitrary function *s*(*i*) relative to that for *s*(*i*) = *s*_*p*_,

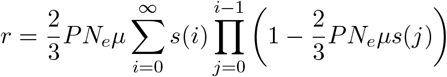

Let,

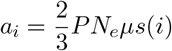

then,

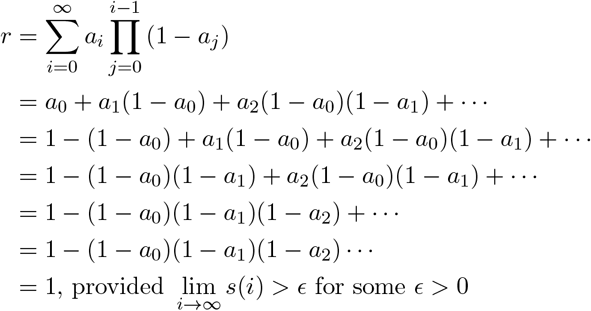

Thus, provided the beneficial allele is always strong, gradual emergence of latent sites does not alter the formula for the fitness effect of the latent load, log 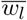.

It is possible that gradual emergence may alter *f*_*l*_, the proportion of new positively selected sites that are latent sites. However, equation 19 shows that *f*_*l*_ is a decreasing function of *s*_*p*_, and thus gradual emergence will tend to push *f*_*l*_ even closer to 1.

## Appendix F: The substitutional load

Assume *PN*_*e*_*s*_*p*_ ≫ 1. Also assume that the selection coefficient of the beneficial allele does not appreciably change between the time it is established and the time it is fixed.

Kimura has determined the expected number of generations until fixation for a beneficial allele with selective advantage *s*, 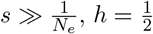, and initial proportion *p*, as approximately (Kimura and Ohta, 1969),

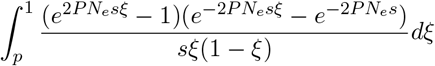

The integrand represents the time (measured in generations) spent at each proportion of the population. A second term dealing with the time spent at proportions less than the initial proportion is negligible and is therefore ignored.

1 − *ξ* is the proportion of the population not containing the beneficial allele. If the log fitness cost of a genomic site not harboring the beneficial allele is *C*, then multiplying the integrand within the integral by *C*(1 − *ξ*) yields the time-aggregated sum of the fitness costs associated with the fixation of the beneficial allele,

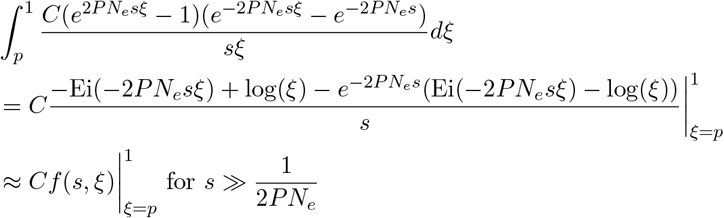

in which,

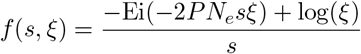

and Ei is the exponential integral,

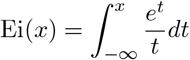

Since *h* is not necessarily equal 0.5 an adjustment to this approach is required. Even with this adjustment, the results are only approximate. For nearly all sexual eukaryotes, the log fitness effect of the substitutional load is far smaller than that of the latent load. Thus this approximation will only produce a small error in the overall genetic load.

Let 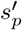 denote the increase in fitness associated with a homozygous beneficial allele relative to a heterozygous beneficial allele. The log fitness of a homozygous beneficial allele is approximately 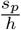, and that of a heterozygous beneficial allele is approximately *s*_*p*_, so,

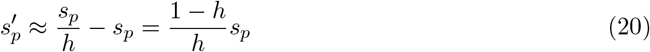

Fixation can be modeled as involving two sequential time intervals, a heuristic approximation adopted for analytical tractability. The first interval covers frequencies from *p* to 50% beneficial allele frequency, with *s* = *s*_*p*_. The second covers frequencies from 50% to 100% beneficial allele frequency, with 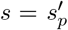. Ignoring rare cases in which organisms are homozygous for the beneficial allele during the first interval, and homozygous for the non-beneficial allele during the second interval, the aggregate log fitness loss associated with the fixation of a beneficial allele, *A*_*s*_, can be approximated in two parts. A cost of 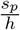 times *f* (*s*_*p*_, *ξ*) with *ξ* ranging from *p* to 0.5. And a cost of 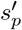 times 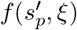 for *ξ* ranging from 0.5 to 1,

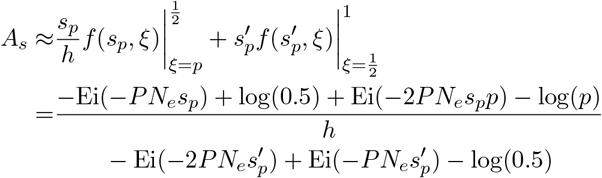

Now, Ei(*x*) ≈ 0 for *x* ≪ −1. Hence, assuming *s*_*p*_ and 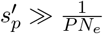,

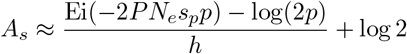

In some cases, a simplification is possible. Ei(*x*) ≈ *γ* + log(|*x*|) for |*x*| ≪ 1 where *γ* is Euler’s constant (0.5772…). It is not guaranteed, but if 2*PN*_*e*_*s*_*p*_*p* ≪ 1, then,

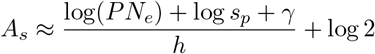

The simplifying condition is the same as the condition for a previously neutral site that is now a positively selected site experiencing a latent load. It thus corresponds to the case in which fixation derives from a single initial mutation. If fixation derives from multiple initial copies of the beneficial allele, *A*_*s*_ will be less than given above.

In the general case, for strong selection, the mean aggregate log fitness loss, 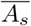, is,

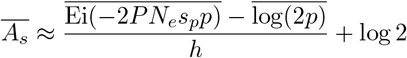

Let *k*_*s*_ ≤ 1 denote a factor reflecting the presence of preexisting established beneficial alleles at new positively selected sites. Then,

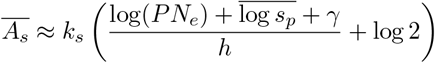

This equation applies under strong selection.

For non-strong selection, a bound on *A*_*s*_ is obtained by combining the maximum fitness loss,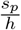, with the maximum mean number of generations to fixation, 2*PN*_*e*_ (Kimura and Ohta, 1969),

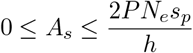

In both cases, the log of mean fitness associated with the substitutional load, log 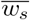, is given by,

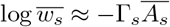

## Appendix G: The genetic load of nearly neutral sites

The analysis of the mutational load thus far applies only when selection dominates over random drift. When |*PN*_*e*_*s*| ≈ 1, selection and drift are in balance. This is the range of nearly neutral sites (Ohta, 1992).

Kimura et al. investigated the mutational load in regimes where selection does not necessarily dominate over drift (Kimura *et al*., 1963), and provided formulas for the total mutational load when |*PN*_*e*_*s*_*n*_| is small or large, but not for the intermediate range. For |2*N*_*e*_*s*_*n*_| *<* 5 in a diploid population with 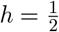, it was found that,

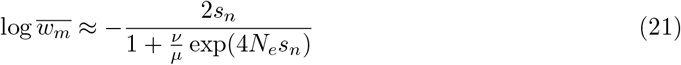

where *ν* is the per site rate of back mutation.

The analysis here extends this finding by deriving a formula for the total mutational load across the full range of selection strengths, and importantly by partitioning the load into a *µ*-responsive and a non-*µ*-responsive component.

Due to random drift, nearly neutral sites can become fixed in the non-beneficial state. Sites that do this might be said to “flap”, spending long periods fixed in both beneficial and deleterious states. The genetic load that results from genetic drift is independent of the mutation rate and is termed the drift load.

Even sites that are not neutral or nearly neutral may spend some time fixed in a deleterious state (Kimura, 1962). Such sites will be termed intermediate sites. It is convenient to treat the genetic load arising from such sites as comprising an effective drift load and effective mutational load.

Assume 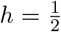. Suppose there is a single base, base *a*, that has a selection coefficient of zero, and each of the other bases has a selection coefficient of −*s*; *s* may be positive or negative, with |*s*| ≪ 1. The mutation rate for base *a* is *µ*, and the aggregate rate of back mutation from the other bases to base *a* is *ν*. This is a simplified model adopted for tractability, ignoring non-additive dominance and multi-allele dynamics.

The site will oscillate between state a, in which it was most recently fixed at base *a*, and state b, in which it was most recently fixed at some other base. Let *t*_*a*_ be the mean time (measured as the number of generations) spent in state a, and *t*_*b*_ be the mean time spent in state b. Let *r*_*a*_ denote the rate of mutation from base a to some other base, and *r*_*b*_ denote the rate of mutation from other bases to base a. Let *A*_*nn*_(−*s*) *<* 0 denote the mean time-aggregated fitness associated with a mutation that fails to fix when in state a, and *A*_*nn*_(*s*) *>* 0 denote the mean time-aggregated fitness gain associated with a mutation that fails to fix when in state b. The total time-averaged log fitness associated with a site, 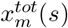, is then given by,

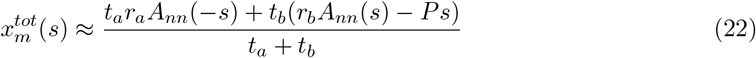

The term −*Ps* occurs in the numerator because it is the fitness effect for state b in the absence of the beneficial allele.

For state a,

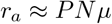

Let *P*_*a*_ denote the probability that a single mutation eventually fixes when the site is in state a. Since initially 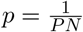, and *s* ≪ 1, from equation 1,

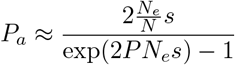

And,

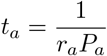

Similarly, let *P*_*b*_ denote the probability that a single mutation eventually fixes when the site is in state b,

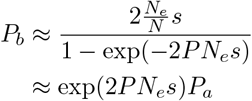

And,

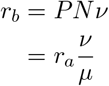

Hence,

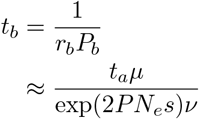

Substituting into equation 22,

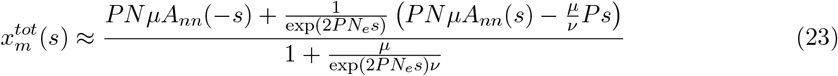

*A*_*nn*_(*s*) can be determined in a manner similar to the substitutional load. Kimura has determined the expected number of generations until extinction of an allele with selective advantage *s* and proportion *p* as (Kimura and Ohta, 1969),

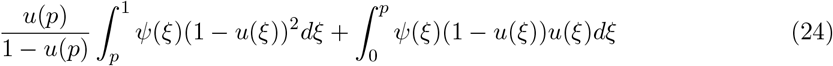

where, using Kimura’s notation,

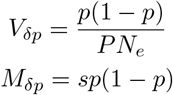

so that,

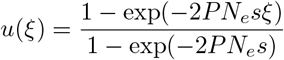

and,

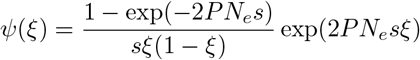

In equation 24, *ξ* is the proportion of haploid genomes with the allele. Multiplying the integrands by *Psξ* therefore gives the time-aggregated fitness,

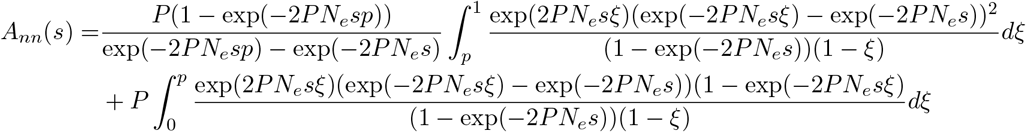

Assuming *N* ≫ 1, since 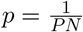, *p* will be small and the second term is negligible. Replacing *ξ* by 1 − *ξ* in the first integral then gives,

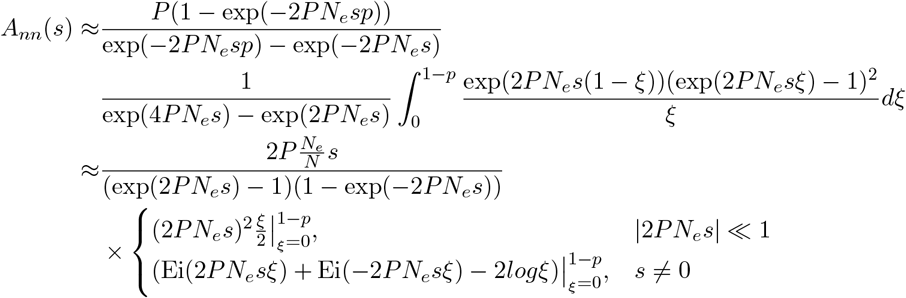

Now Ei(*x*) ≈ *γ* + log |*x*| when |*x*| ≪ 1, Ei(*x*) ≈ 0 when *x* ≪ −1, and 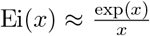 when *x* ≫ 1, so,

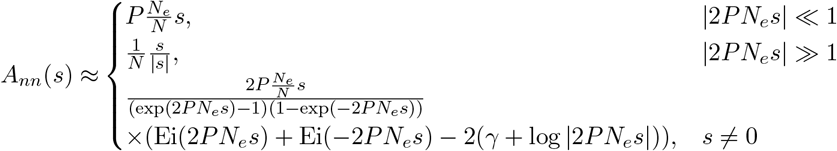

From this it follows that *A*_*nn*_(−*s*) ≈ −*A*_*nn*_(*s*). Hence, by equation 23,

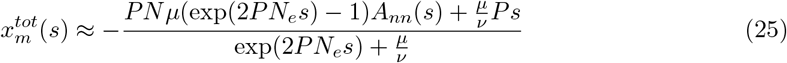

Assuming the Jukes-Cantor model 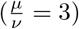, in several special cases this gives,

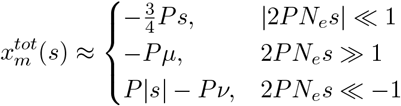

which agrees with intuition and with equation 14.

For the general case, with a Jukes-Cantor model, Figure 3 shows the two components of 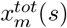; a *µ*-responsive component and a non-*µ*-responsive component (for fixed 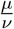), which are summed to give 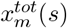. See S1 File for the plotting code. In the figure the fitness effect of the *µ* component is divided by *Pµ*, and the non-*µ* component is multiplied by *N*_*e*_. The non-*µ* component starts at zero, reaches a maximum of around 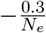at *PN*_*e*_*s* ≈ 0.8, and then slowly falls back to almost zero by the time *PN*_*e*_*s* = 4. The *µ* component starts at zero, interestingly increases to a maximum value larger than −*Pµ* at *PN*_*e*_*s* ≈ 2.9, and then asymptotically falls back to the value −*Pµ*.

**Figure 3:**
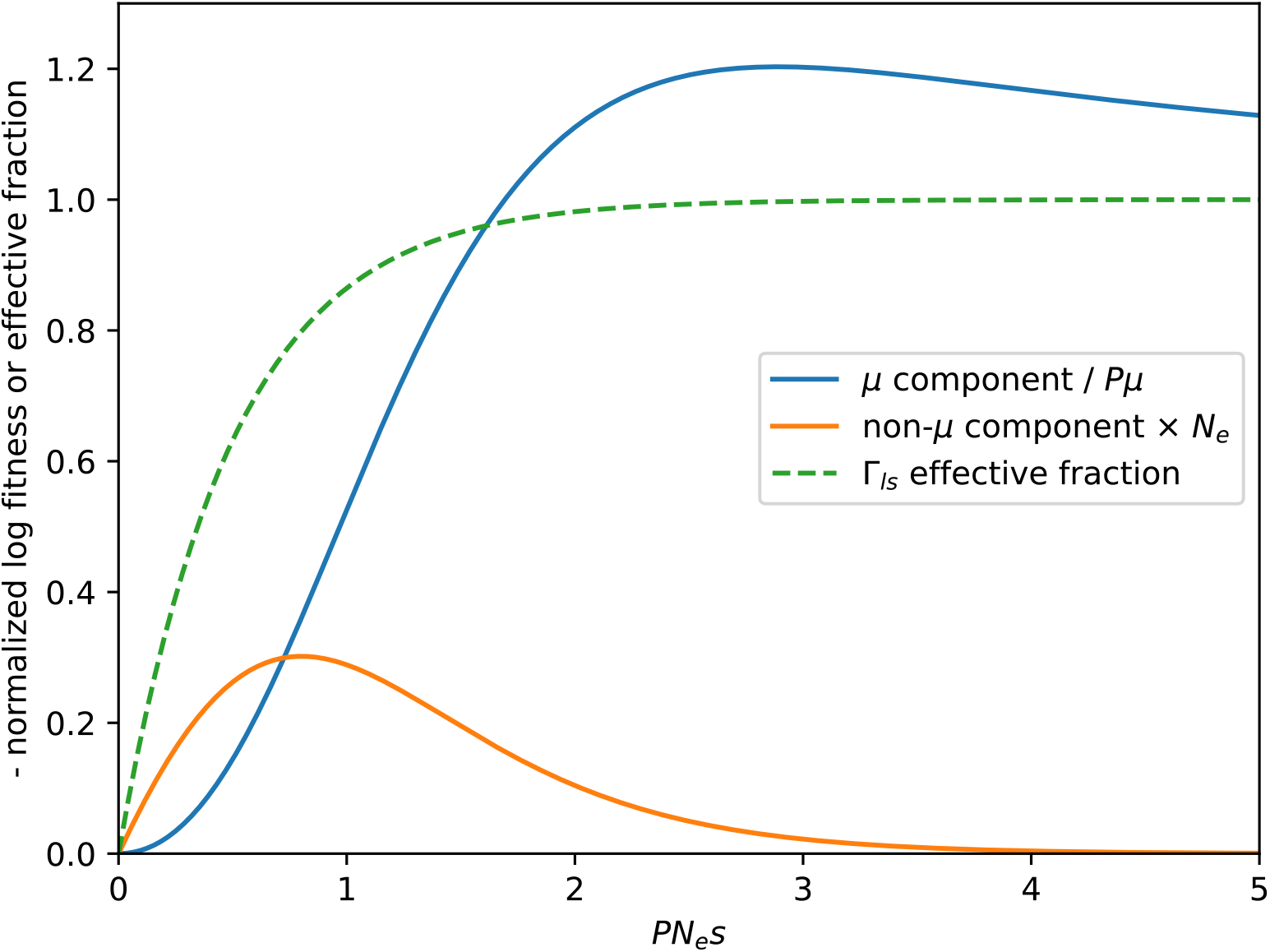
Time-averaged log fitness components for a site. Ploidy *P*, effective population size *N*_*e*_, selection coefficient *s*, mutation rate *µ*, and coefficient of dominance, 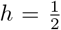, assuming a Jukes-Cantor model. Also shown is the Γ_*ls*_ effective fraction.

Also shown for reference in Figure 3 is the Γ_*ls*_ effective fraction, 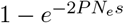. For small *PN*_*e*_*s*, the Γ_*ls*_ effective fraction falls off to a lesser extent than the *µ* component.

Figure 3 shows that the non-*µ* component of a nearly neutral site can be substantially larger than the standard *Pµ* associated with the mutational load when *PN*_*e*_*µ* ≪ 1. Consequently, depending on the number of nearly neutral sites, and their selection coefficients, the fitness loss associated with them could be substantial. Such sites are nearly neutral with respect to individual selection, but not necessarily with respect to aggregate fitness.

An illustrative example typical of vertebrates helps clarify this point. Consider an exponential distribution of fitness effects for new positively selected sites with mean effect size 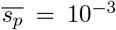. Numerical integration of the non-*µ* component of equation 25 weighted by the distribution of fitness effects for a Jukes-Cantor model with *P* = 2 and *N*_*e*_ = 10^5^ yields a log of mean fitness of −2.4*×*10^−8^ per site. See S1 File for the numerical integration code. This is the mean value of the non-*µ* component of nearly neutral sites. Meanwhile, the log of mean fitness from the mutational load of those new sites that will become conserved is approximately −*Pµ*. All non-neutral sites were once new positively selected sites. Therefore, assuming their selection coefficients have remained unchanged, for *µ* = 10^−8^ the drift load would be larger than the mutational load.

Let *D*_*nn*_(*s*) denote the genomic density of nearly neutral sites with selection coefficient *s* for some definition of nearly neutral. Then the log fitness loss associated with nearly neutral sites, log 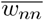, is given by,

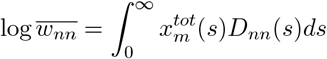

At times it may be useful to decompose log 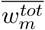 into its *µ* and non-*µ* responsive components, and speak of an effective mutational load having fitness effect log 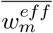 which combines the *µ* component of nearly neutral sites with the mutational load of conserved sites, and to speak of the drift load having fitness effect log 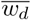 which consists of just the non-*µ* component of nearly neutral and intermediate sites.

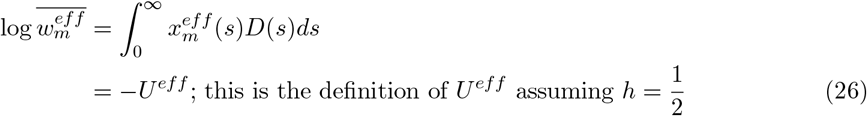

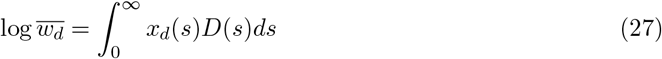

where *D*(*s*) is the density of sites with selection coefficient *s* in the genome, and for 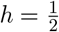,

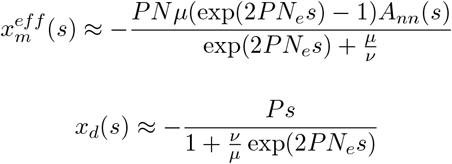

This last equation is in agreement with equation 21 derived by Kimura et al. (Kimura *et al*., 1963).

This analysis was for 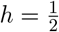. Other dominance coefficients may quantitatively alter the results, but do not appear likely to prevent the decomposition.

## Data availability

All relevant data are within the manuscript and its Supporting information.

## Supporting information

S1 File - Nearly neutral log fitness software and results.

https://doi.org/10.5281/zenodo.18488042

## Acknowledgments

I am grateful to Michael Lynch for the time he spent reviewing this manuscript and for providing valuable feedback. I am also grateful to Steven Greidinger, Philip Gerrish, and Benjamin Good for the time they spent reviewing an earlier related manuscript.

## Conflict of interest statement

The author declares that they have no conflicts of interest in relation to the content of this manuscript.

## Notes

https://doi.org/10.5281/zenodo.13901486

### Competing Interest Statement

The authors have declared no competing interest.

### Summary of Updates

Change focus of the manuscript to the latent load.

